# De Novo Assembly of a Chromosome-Scale Reference Genome for the Northern Flicker *Colaptes auratus*

**DOI:** 10.1101/2020.08.19.257683

**Authors:** Jack P. Hruska, Joseph D. Manthey

**Affiliations:** Department of Biological Sciences, Texas Tech University. Lubbock, Texas, USA

**Keywords:** *Colaptes auratus*, woodpeckers, PacBio, HiC, genome assembly

## Abstract

The northern flicker, *Colaptes auratus*, is a widely distributed North American woodpecker and a long-standing focal species for the study of ecology, behavior, phenotypic differentiation, and hybridization. We present here a highly contiguous *de novo* genome assembly of *C. auratus,* the first such assembly for the species and the first published chromosome-level assembly for woodpeckers (Picidae). The assembly was generated using a combination of short-read Chromium 10x and long-read PacBio sequencing, and further scaffolded with chromatin conformation capture (Hi-C) reads. The resulting genome assembly is 1.378 Gb in size, with a scaffold N50 of 43.948 Mb and a scaffold L50 of 11. This assembly contains 87.4 % - 91.7 % of genes present across four sets of universal single-copy orthologs found in tetrapods and birds. We annotated the assembly both for genes and repetitive content, identifying 18,745 genes and a prevalence of ~ 28.0 % repetitive elements. Lastly, we used four-fold degenerate sites from neutrally evolving genes to estimate a mutation rate for *C. auratus*, which we estimated to be 4.007 × 10^−9^ substitutions / site / year, about 1.5x times faster than an earlier mutation rate estimate of the family. The highly contiguous assembly and annotations we report will serve as a resource for future studies on the genomics of *C. auratus* and comparative evolution of woodpeckers.

## INTRODUCTION

The northern flicker *Colaptes auratus* is a polytypic North America woodpecker with a distribution spanning from Alaska to northern Nicaragua, Cuba, and the Cayman Islands. *C. auratus* consists of up to 13 described subspecies (Gill, Donsker, and Rasmussen, 2020) and five morphological groups (Short 1982). Currently, the taxonomy of *C. auratus* is uncertain; some authorities consider it to form a species complex along with the gilded flicker *Colaptes chrysoides*, while others have suggested that one of the subspecies, *C.a. mexicanoides*, is best considered a separate species (del Hoyo et al., 2014). In addition, hybridization between morphological groups in secondary contact is prevalent, primarily between the yellow-shafted and red-shafted flickers, who form a hybrid zone that extends from northern Texas to southern Alaska (K. L. Wiebe and W.S. Moore 2020). The yellow-shafted/red-shafted hybrid zone has become a prominent study system for the consequences of secondary contact (e.g., Moore and Koenig 1986; Karen L. Wiebe 2000). Despite there being marked phenotypic differentiation between red-shafted and yellow-shafted flickers, genetic divergence between these groups is remarkably shallow, even when sampling thousands of markers across the genome (Manthey et al. 2015; Aguillon et al. 2018). The paradoxical conjunction of shallow genetic divergence and marked phenotypic differentiation echoes the genomic dynamics of other avian hybrid zones, namely the golden-winged *Vermivora chrysoptera* and blue-winged *Vermivora cyanoptera* complex, wherein only a few genomic regions associated with genes that determine plumage color and pattern differentiate the two species (Toews et al. 2016). A chromosome-level reference genome for the complex will not only facilitate the identification of the genetic basis of phenotypes (Kratochwil and Meyer 2015), a long-standing goal in evolutionary biology research, but also provide researchers a valuable resource for the examination of emerging fields in genome biology, such as the evolutionary dynamics of transposable element proliferation (Manthey, Moyle, and Boissinot 2018), for which woodpeckers are especially well suited.

Here, we describe Caur_TTU_1.0, a *de novo* assembly that was built from a wild caught *C. auratus* female. We used three sequencing strategies: 10x Chromium, PacBio, and chromatin conformation capture (Hi-C) to assemble the first published chromosome level genome for *C. auratus* and Picidae. As whole genome sequencing becomes more feasible and prevalent, high-quality reference genomes will undoubtedly serve as essential resources. We expect the chromosome-level assembly presented here will be of great use to those interested in the genomic evolution of woodpeckers and birds, at large.

## METHODS AND MATERIALS

### DNA extraction, Library Preparation, and Sequencing

We obtained breast muscle tissue from a vouchered *C. auratus* specimen (MSB 48083) deposited at the Museum of Southwestern Biology (MSB). The specimen was a wild female collected on 11 July 2017 in Cibola County, New Mexico (see MSB database for complete specimen details). We used a combination of 10x Chromium, PacBio, and Hi-C sequencing data for genome assembly. 10x Chromium library sequencing was carried out by the HudsonAlpha Institute for Biotechnology (Huntsville, AL, USA). They performed high molecular weight DNA isolation, quality control, library preparation, and shotgun sequencing on one lane of an Illumina HiSeqX. For long-read PacBio sequencing, we used the services of RTL Genomics (Lubbock, TX, USA). They performed high molecular weight DNA isolation using Qiagen (Hilden, Germany) high-molecular weight DNA extraction kits, PacBio SMRTbell library preparation, size selection using a Blue Pippin (Sage Science), and sequencing on six Pacific Biosciences Sequel SMRTcells 1M v2 with Sequencing 2.1 reagents. Hi-C library preparation was performed with an Arima Genomics Hi-C kit (San Diego, CA, USA) by the Texas A&M University Core facility. The Hi-C library was then sequenced on a partial lane of an Illumina NovaSeq S1 flow cell at the Texas Tech University Center for Biotechnology and Genomics.

### Genome Assembly, Polishing, Scaffolding and Quality Assessment

We generated an initial assembly using the raw PacBio long reads with CANU v 1.7.1 (Koren et al. 2017). Reads were corrected, trimmed, and assembled using CANU default parameters, while specifying a normal coarse sensitivity level (- corMhapSensitivity flag), setting the expected fraction error in an alignment of two corrected reads to 0.065 (- correctedErrorRate flag) and setting the estimated genome size to 1.6 Gb, which corresponds with previous estimates within *Colaptes* (Wright, Gregory, and Witt 2014). We subsequently polished the PacBio assembly using the 10x Chromium sequencing reads with one iteration of the PILON v 1.22 (Walker et al. 2014) pipeline, which consisted of several steps. We first used bbduk, part of the BBMap v38.22 package (Bushnell 2014), to trim adapters and quality filter the raw 10x Chromium reads. We then used the BWA-MEM implementation of the Burrows-Wheeler algorithm in BWA v 0.7.17 (Li and Durbin 2010) to align these filtered reads to the PacBio assembly. We used samtools v 1.9 (Li et al. 2009) to sort and index the resulting BAM file, which along with the PacBio assembly, was input to PILON. Following polishing, we then performed scaffolding of the PacBio assembly with the 10x Chromium reads using ARCS (Yeo et al. 2018). An interleaved linked reads file of the 10x Chromium reads produced in LongRanger v 2.2.2 was subsequently input to the ARCS pipeline, which implements LINKS v1.8.5 (Warren et al. 2015). Three rounds of ARCS were performed, wherein each round multiple iterations of the pipeline were run to evaluate which parameter combination produced the assembly of highest quality.

Default parameters of the pipeline were used, with the following exceptions: (1) the link ratio between two best contig pairs (- a flag), which was set to 0.5; (2) the minimum link number of links to compute scaffold (- l flag), which was set to 3; (3) the minimum sequence identify (- s flag), was varied between 97,98, and 99; (4) the contig head/tail length for masking alignments was varied between 10k, 30k, 60k, and 100k. After all iterations were run, the assembly with greatest scaffold N50 and size was selected and used in subsequent rounds. Lastly, we used the Hi-C reads to further scaffold and fix mis-assemblies using the 3D-DNA pipeline (Dudchenko et al. 2017; Durand et al. 2016).

To assess the spatial order of the scaffolds of the Caur_TTU_1.0 assembly, we aligned it to the Chicken *Gallus gallus* chromosome-level assembly (GRCg6a, GCF_000002315.6, https://www.ncbi.nlm.nih.gov/genome/?term=Gallus) using the nucmer module of MUMMER v 4.0.0b2 (Kurtz et al. 2004). We subsequently filtered alignments using MUMMER’s delta-filter module while setting the minimum alignment identity to 70% and allowing many-to-many alignments. A tab-delimited text file that include information on the position, percent identity, and length of each alignment was produced using MUMMER’s show-coords module. This file was used as input to create a synteny plot with OmicCircos (Hu et al. 2014; R Core Team 2018). Subsequently, the Caur_TTU_1.0 scaffolds were renamed according to their corresponding Chicken chromosome. Scaffolds that did not show strong synteny to Chicken chromosomes were not renamed.

Genome assembly metrics were obtained using the function stats.sh from the BBMap v 38.22 package (Bushnell 2014). Genome completeness was estimated using Tetrapoda and Aves single-copy orthologous gene sets from both BUSCO v3 (Simão et al. 2015; Waterhouse et al. 2018) and BUSCO v4 (Seppey, Manni, and Zdobnov 2019). We submitted our genome assembly to the NCBI genome submission portal, where a scan for contaminants detected no abnormalities in our assembly.

### Genome annotation

#### Repetitive element annotation and window analysis

We annotated transposable elements (TEs) and repetitive content in the Caur_TTU_1.0 assembly using a custom *de novo* repeat library and RepBase vertebrate database v 24.03 (Jurka et al. 2005). The custom repeat library was constructed from the *C. auratus* genome assembly (prior to Hi-C scaffolding) and other in-progress lab genome assembly projects in songbirds (File S15).

Using the RepBase vertebrate database and the *de novo* repeat library, we used RepeatMasker v 1.332 (A. Smit, Hubley, and Green 2015) to mask and summarize repetitive and transposable elements in the Caur_TTU_1.0 assembly (Files S16 & S17). An interspersed repeat landscape was then produced for the Caur_TTU_1.0 assembly using the RepeatMasker scripts calcDivergenceFromAlign.pl and createRepeatLandscape.pl. The spatial distribution of repetitive content across the Caur_TTU_1.0 assembly was evaluated using custom R scripts (R Core Team 2018), first by removing overlapping elements from the RepeatMasker output, followed by a calculation of repetitive element content of the Chicken-renamed scaffolds across 100 kbp non-overlapping sliding windows.

To generate the custom repeat library, we first input the *C. auratus* assembly that lacked Hi-C scaffolding to RepeatModeler v 1.10.11 (A. F. Smit and Hubley 2008) to identify repeats *de novo*. RepeatModeler identifies repeats according to homology, repeats and repetitiveness with the programs RECON (Bao and Eddy 2002), RepeatScout (Price, Jones, and Pevzner 2005) and Tandem Repeats Finder (Benson 1999). We then removed RepeatModeler sequences that were ≥ 98 % identical to the RepBase vertebrate database. Next, we used blastn v 2.9.0 (Camacho et al. 2009) and bedtools v 2.29.2 (Quinlan and Hall 2010) to extract sequence matches to these novel repeats from the aforementioned assembly. We then used these sequences to create consensus sequences for each novel repetitive element using the following workflow: (1) alignment of reads using MAFFT (Katoh and Standley 2013) as implemented in Geneious (BioMatters Ltd.); (2) generation of 50% majority consensus sequences from these alignments in Geneious; (3) trimming ambiguous nucleotides on the ends of consensus sequences. For novel repetitive elements whose ends were not recovered in the generation of the consensus sequences, we repeated the prior procedure and extracted sequences from the reference genome with 1000 bp flanks on each side of the blastn match, followed by alignment and consensus sequence generation as mentioned above (Platt, Blanco-Berdugo, and Ray 2016). This process was repeated up to three times. We then BLASTed all novel repeats against the RepBase database to assess similarity via homology to previously characterized elements. Similarity to RepBase elements was used for naming purposes.

#### Gene annotation and window analysis

We employed MAKER v 2.31.10 (Cantarel et al. 2008) to annotate putative genes in the Caur_TTU_1.0 assembly. We used the custom repeat library and protein datasets of four species in MAKER to predict genes. The species included were: (1) *Picoides pubescens* (GCF_000699005.1_ASM69900v1_Picoides_pubescens_protein.faa), (2) *Merops nubicus* (GCF_000691845.1_ASM69184v1_Merops_nubicus_protein.faa), (3)

#### Apaloderma vittatum

(GCF_000703405.1_ASM70340v1_Apaloderma_vittatum.protein.faa), (4) and *Buceros rhinoceros* (GCF_000710305.1_ASM71030v1_Buceros_rhinoceros_protein.faa) (Zhang et al. 2014). We then used these predictions to train the *ab initio* gene predictors SNAP (Korf 2004) and Augustus v.3.2.3 (Stanke et al. 2008). Lastly, using the SNAP and Augustus-trained gene models, we ran a second round of MAKER to annotate genes in the Caur_TTU_1.0 assembly. The spatial distribution of coding sequences (CDS) across the Chicken-renamed scaffolds of the Caur_TTU_1.0 assembly was evaluated using a custom R script (R Core Team 2018).

#### Mutation rate estimation

We extracted the putative coding sequences (CDS) (File S14) from the Caur_TTU_1.0 assembly using the final MAKER output and bedtools. In addition, we downloaded the CDS for *Apaloderma vittatum*, *Merops nubicus*, and *Buceros rhinoceros* for homology-based comparisons (using the same genomes containing the aforementioned protein datasets). We performed a reciprocal BLAST of all species versus *C. auratus* using blastn to identify putative homologues across all four species.

To put the evolution of the CDS regions in a timed evolutionary context, we downloaded a phylogenetic tree comprising all orders of Neoaves (Jarvis et al. 2014) and pruned the tree to the four representative orders covered by our CDS downloads and the Caur_TTU_1.0 assembly using the R package ape (Paradis, Claude, and Strimmer 2004): Piciformes, Coraciiformes, Trogoniformes, and Bucerotiformes.

We used T-Coffee (Notredame, Higgins, and Heringa 2000) to align the putative homologues between the four passerine species. T-Coffee translates nucleotide sequences, aligns them using several alignment algorithms, takes the averaged best alignment of all alignments, and back translates the protein alignments to provide a nucleotide alignment for each gene. Prior to back-translating, we removed any gaps in the protein alignments using trimAl (Capella-Gutiérrez, Silla-Martínez, and Gabaldón 2009).

With the alignments for all genes, we tested for selection using the gene-wide and branch-specific tests for selection utilized in CODEML (Yang 1997). Any alignments with gene-wide or branch-specific evidence for selection were removed for mutation-rate analyses, after correcting for multiple tests using the Benjamini and Hochberg (Benjamini and Hochberg 1995) method to control false discovery rate. From each gene alignment, we used the R packages rphast, Biostrings, and seqinr (Hubisz, Pollard, and Siepel 2011; Pagès et al. 2017; Charif et al. 2007) to extract four-fold degenerate sites from each alignment. We concatenated the four-fold degenerate sites (N ~ 528 thousand) and used jModelTest2 (Darriba et al. 2012) to determine an appropriate model of sequence evolution. We used the GTR + I model of sequence evolution in PhyML v 3.3.20190321 (Guindon and Gascuel 2003; Guindon et al. 2010)and a user-specified tree (from Jarvis et al. 2014) to estimate branch lengths based on the four-fold degenerate sites. Lastly, we used the *Colaptes*-specific branch length of this tree along with divergence time estimates (also from Jarvis et al. 2014) to estimate a mean and 95% HPD distribution of potential *Colaptes-*lineage specific mutation rates.

## RESULTS AND DISCUSSION

### Sequencing, genome assembly, and synteny mapping

Reads were generated across three sequencing approaches, including 3.94 × 10^6^ Pacific Biosciences (PacBio) long-reads (~34x coverage), 4.47 × 10^8^ 10x Chromium paired end reads (~58x raw coverage), and 3.25 × 10^8^ Hi-C paired end reads (> 24,000x physical distance coverage after deduplication). The final assembly had an L50 of 11 scaffolds and an N50 of 43.938 Mbp (Figure 1A; Table 1). In terms of contiguity (L50 and N50), this assembly represents a ~3x improvement over a recently published long-read based Picidae assembly (*Melanerpes aurifrons* GCA_011125475.1; Wiley and Miller 2020) and represents the first published chromosome-level assembly for Piciformes. BUSCO results also suggested this assembly is of high-quality, with modestly high recovery of complete bird-specific and tetrapod-specific gene groups (87.4 % - 91.7 %; Table 2; Files S1-S12). While a higher gene group recovery rate would be expected for a highly-contiguous assembly, we highlight that these results correspond with studies that have found that greater assembly contiguity often does not result in an increased gene group recovery rate, and if an increase is noted, it is often modest (Korlach et al. 2017; Low et al. 2019). Indeed, we find our recovery rates to be similar to those of the *Melanerpes aurifrons* assembly, with 92.6 % of complete BUSCO gene groups recovered from the aves_odb9 dataset (Wiley and Miller 2020). We recovered a high degree of one-to-one synteny with the Chicken *Gallus gallus* chromosomes, particularly between those of small and medium size (Figure 1B). However, we note that one-to-one synteny to the *Gallus* assembly was lacking for the larger chromosomes, indicative of chromosomal splitting since the *Gallus*-*Colaptes* common ancestor has occurred. Members of Picidae are known for containing a high number of chromosomes, particularly micro-chromosomes (Kaul and Ansari 1978), and karyotypes of *Colaptes* have been shown to consistently have a larger number of chromosomes when compared to *Gallus* (Pollock and Fechheimer 1976; de Oliveira et al. 2017).

**Table 1.** Genome Assembly Metrics calculated using BBMap.

| Statistic | Caur_TTU_1.0 |
| --- | --- |
| # scaffolds / contigs | 2369 / 9565 |
| Largest scaffold / contig | 117.313 Mbp / 15.844 Mbp |
| Total Length | 1.378 Gbp |
| Scaffold / contig N50 | 43.948 Mbp / 826.96 Kbp |
| Scaffold / contig N90 | 14.604 Mbp / 50.09 Kbp |
| Scaffold / contig L50 | 11 / 281 |
| Scaffold / contig L90 | 33 / 4370 |
| GC (%) | 44.93 |

**Table 2.** BUSCO output using tetrapoda_odb9, tetrapoda_odb10, aves_od9 and aves_odb10 databases.

|  | tetrapoda_odb9 | tetrapoda_odb10 | aves_odb9 | aves_odb10 |
| --- | --- | --- | --- | --- |
| **Complete BUSCOs** | 3623 (91.7%) | 4670 (87.9%) | 4416 (89.9%) | 7294 (87.4%) |
| **Complete and single-copy BUSCOs** | 3594 (91.0%) | 4617 (86.9%) | 4342 (88.3 %) | 7224 (86.6%) |
| **Complete and duplicated BUSCOs** | 29 (0.7%) | 53 (1.0%) | 74 (1.5 %) | 70 (0.8%) |
| **Fragmented BUSCOs** | 147 (3.7%) | 124 (2.3%) | 227 (4.6 %) | 219 (2.6%) |
| **Missing BUSCOs** | 180 (4.6 %) | 516 (9.8%) | 272 (5.6 %) | 825 (10.0%) |
| **Total BUSCO groups searched** | 3950 | 5310 | 4915 | 8338 |

**Figure 1.**
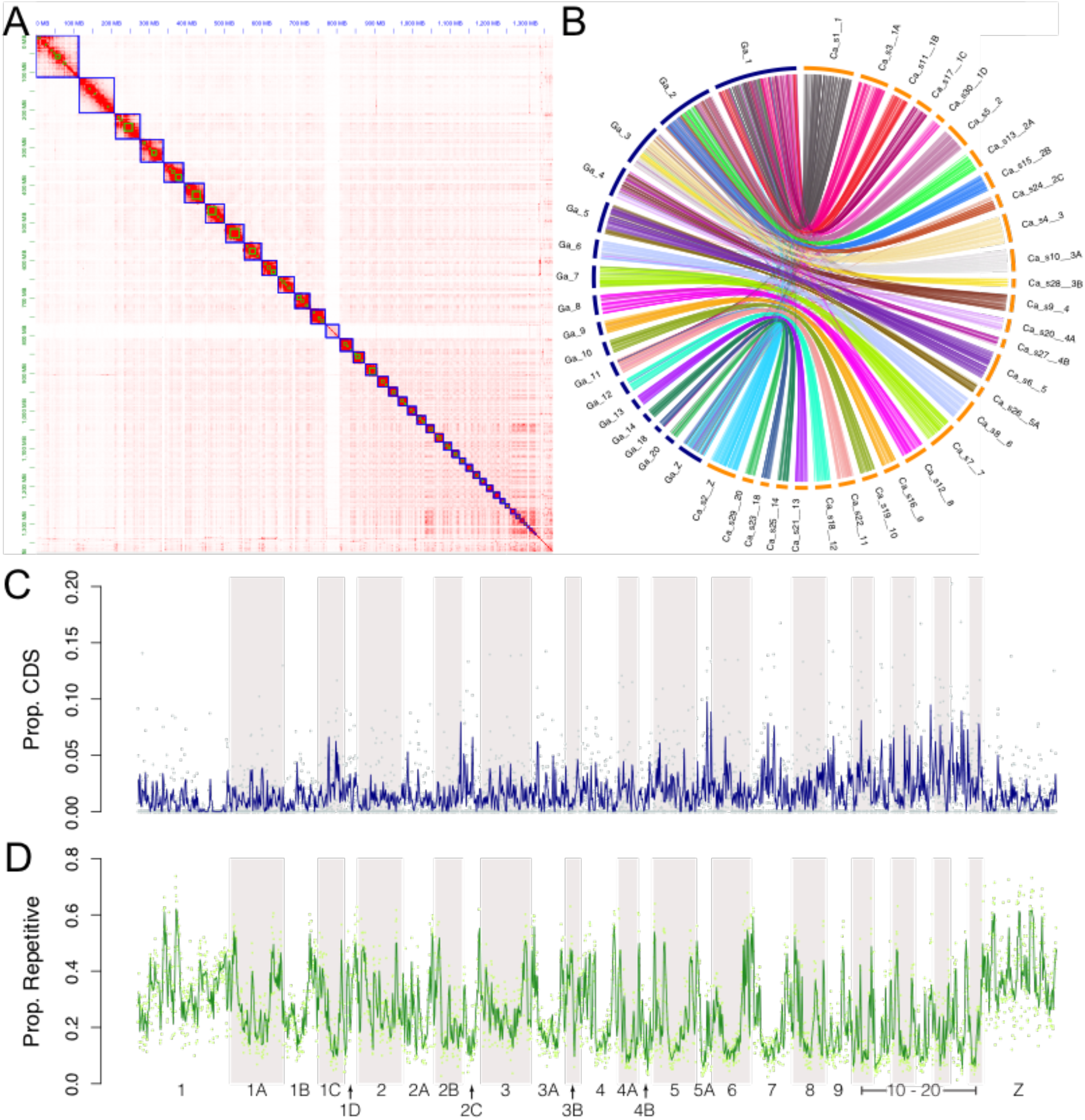
Characteristics of the Caur_TTU_1.0 assembly **(A)** Hi-C scaffolding contact map. Relative contact between contigs is indicated by the intensity of red. Blue squares indicate scaffold boundaries. **(B)** Synteny map of Caur_TTU_1.0 (right; orange) scaffolds to *Gallus gallus* chromosomes (left; blue). **(C)** Proportions of coding sequence (CDS; top panel) and repetitive elements (bottom panel) across 100 kbp sliding non-overlapping windows of the Chicken-aligned Caur_TTU_1.0 scaffolds. Lines indicate mean values across ten sliding non-overlapping windows.

### Genome annotation, window analysis, and mutation rate estimation

Repetitive elements make up a large portion of Caur_TTU_1.0, comprising ~ 386 Mb (~28 %) of the assembly. The presence of the retrotransposon superfamily CR1 (chicken repeat 1) was particularly prevalent, comprising ~ 20.9% (~ 287 Mbp) of the assembly. Two independent waves of CR1 proliferation were detected, with a large proportion of CR1 elements being of relatively young or medium age, as estimated by a molecular clock (Figure 2). These results echo Manthey et al. (2018), who also uncovered multiple independent waves of CR1 proliferation across Piciformes. Window analysis of repetitive elements suggested that the distribution of these elements was uneven across the assembly, both within and across scaffolds (Figure 1C). Repetitive element content was particularly prevalent near scaffold boundaries and on the Z chromosome, with local repetitive densities reaching ~ 60%. High repetitive content on the Z chromosome has been previously reported in birds (Kapusta and Suh 2017).

**Figure 2.**
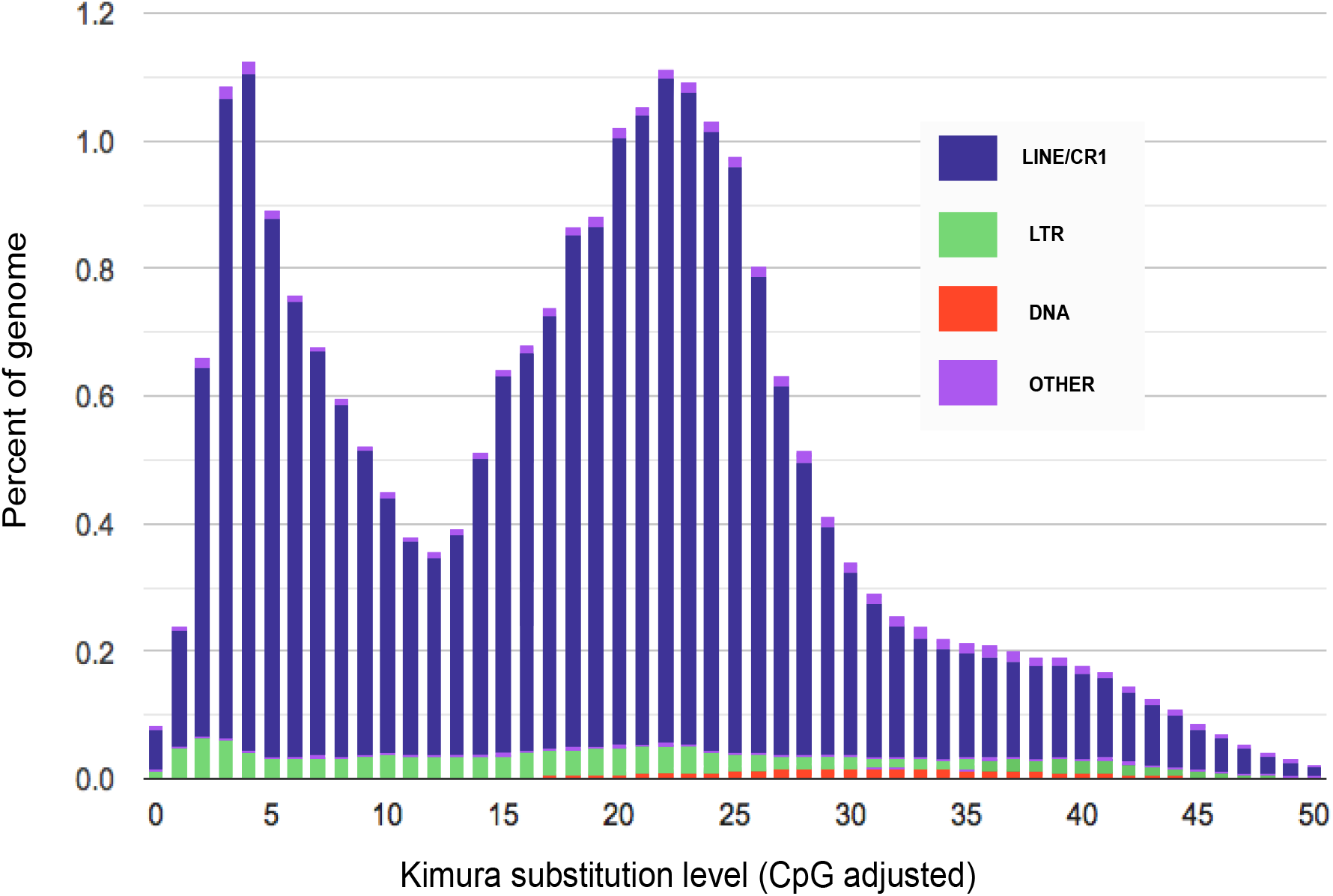
Caur_TTU_1.0 divergence landscape of transposable element classes. Relative abundance and age of each class is shown.

Two rounds of the MAKER annotation pipeline identified a total of 18,745 genes (mean length: 14,676.1 bp) and 149,433 exons (mean length: 161.351 bp) (File S13). The quantity of genes and exons recovered is in line with previously annotated bird genomes (Zhang et al. 2014). The distribution of CDS across 100 kbp sliding windows of the Caur_TTU_1.0 assembly revealed that, as expected, these sequences comprised a smaller fraction of autosomal and sex chromosomes when compared to repetitive elements (Figure 1C).

The mutation rate analysis of four-fold degenerate sites from neutrally evolving genes suggests that the mean rate in *C. auratus* is 4.007 × 10^−9^ substitutions / site / year; with a 95% HPD = 3.525 × 10^−9^ – 4.976 × 10^−9^. This rate is ~1.5x higher than a previous estimate of the Downy Woodpecker *Dryobates pubescens* (2.42 × 10^−9^; Nadachowska-Brzyska et al. 2015). While these results could be reflecting biologically distinct mutation rates between species of woodpeckers, we also acknowledge this discrepancy in results could result from differing methodological choices. Therefore, we urge caution when interpreting this result.

## DATA AVAILABILITY

The Caur_TTU_1.0 assembly is available at NCBI (BioProject PRJNA616131; Genome JAAWVA000000000). All associated raw sequencing data, PacBio (SRR12364887), Chromium 10x (SRR12363123) and Hi-C (SRR12363461) are available from NCBI SRA. Scripts, associated files, and workflows used for this project are available on GitHub (github.com/jphruska/Colaptes_genome). Outputs from BUSCO, Maker, RepeatMasker along with the custom repeat library are deposited as supplemental files in figshare.

## ACKNOWLEDGEMENTS

We would like to thank Christopher Witt and Andrew Johnson at the Museum of Southwestern Biology for granting access to the tissue sample used in this study. We thank Mohamed Fokar at the Texas Tech Center for Biotechnology and Genomics for assistance with Hi-C sequencing. Sequencing was supported by Texas Tech University start-up funding to JDM. The Texas Tech University High Performance Computing Center supported most of the computational analyses. JPH and JDM both performed analyses and wrote, reviewed, and approved the manuscript.

## LITERATURE CITED

Aguillon, Stepfanie M., Leonardo Campagna, Richard G. Harrison, and Irby J. Lovette. 2018. “A Flicker of Hope: Genomic Data Distinguish Northern Flicker Taxa despite Low Levels of Divergence.” The Auk: Ornithological Advances 135 (3): 748–766.

Bao, Zhirong, and Sean R. Eddy. 2002. “Automated de Novo Identification of Repeat Sequence Families in Sequenced Genomes.” Genome Research 12 (8): 1269–1276.

Benjamini, Yoav, and Yosef Hochberg. 1995. “Controlling the False Discovery Rate: A Practical and Powerful Approach to Multiple Testing.” Journal of the Royal Statistical Society: Series B (Methodological) 57 (1): 289–300.

Benson, Gary. 1999. “Tandem Repeats Finder: A Program to Analyze DNA Sequences.” Nucleic Acids Research 27 (2): 573–580.

Bushnell, Brian. 2014. “BBMap: A Fast, Accurate, Splice-Aware Aligner.” Lawrence Berkeley National Lab.(LBNL), Berkeley, CA (United States).

Camacho, C., G. Coulouris, V. Avagyan, N. Ma, J. Papadopoulos, and K. Bealer. 2009. “BLAST plus: Architecture and Applications. BMC Bioinformatics.” BioMed Central 10 (421): 1.

Cantarel, Brandi L., Ian Korf, Sofia MC Robb, Genis Parra, Eric Ross, Barry Moore, Carson Holt, Alejandro Sánchez Alvarado, and Mark Yandell. 2008. “MAKER: An Easy-to-Use Annotation Pipeline Designed for Emerging Model Organism Genomes.” Genome Research 18 (1): 188–196.

Capella-Gutiérrez, Salvador, José M. Silla-Martínez, and Toni Gabaldón. 2009. “TrimAl: A Tool for Automated Alignment Trimming in Large-Scale Phylogenetic Analyses.” Bioinformatics 25 (15): 1972–1973.

Charif, Delphine, Jean R. Lobry, U. Bastolla, M. Porto, H. E. Roman, and M. Vendruscolo. 2007. “Structural Approaches to Sequence Evolution: Molecules, Networks, Populations.” Biological and Medical Physics, Biomedical Engineering (Ed. Bastolla U, Porto M, Roman HE, Vm)(Springer-Verlag, 2007).

Darriba, Diego, Guillermo L. Taboada, Ramón Doallo, and David Posada. 2012. “JModelTest 2: More Models, New Heuristics and Parallel Computing.” Nature Methods 9 (8): 772–772.

Dudchenko, Olga, Sanjit S. Batra, Arina D. Omer, Sarah K. Nyquist, Marie Hoeger, Neva C. Durand, Muhammad S. Shamim, Ido Machol, Eric S. Lander, and Aviva Presser Aiden. 2017. “De Novo Assembly of the Aedes Aegypti Genome Using Hi-C Yields Chromosome-Length Scaffolds.” Science 356 (6333): 92–95.

Durand, Neva C., James T. Robinson, Muhammad S. Shamim, Ido Machol, Jill P. Mesirov, Eric S. Lander, and Erez Lieberman Aiden. 2016. “Juicebox Provides a Visualization System for Hi-C Contact Maps with Unlimited Zoom.” Cell Systems 3 (1): 99–101.

Gill, F, D Donsker, andP Rasmussen, eds. n.d. “IOC World Bird List (Version 10.1).” 2020. https://doi.org/10.14344/IOC.ML.10.1.

Guindon, Stéphane, Jean-François Dufayard, Vincent Lefort, Maria Anisimova, Wim Hordijk, and Olivier Gascuel. 2010. “New Algorithms and Methods to Estimate Maximum-Likelihood Phylogenies: Assessing the Performance of PhyML 3.0.” Systematic Biology 59 (3): 307–321.

Guindon, Stéphane, and Olivier Gascuel. 2003. “A Simple, Fast, and Accurate Algorithm to Estimate Large Phylogenies by Maximum Likelihood.” Systematic Biology 52 (5): 696–704.

Hoyo, J. del, N.J. Collar, D.A. Christie, A. Eliiott, and L.D.C. Fishpool. 2014. HBW and BirdLife International Illustrated Checklist of the Birds of the World. Barcelona, Spain and Cambridge, UK.: Lynx Edicions BirdLife International.

Hu, Ying, Chunhua Yan, Chih-Hao Hsu, Qing-Rong Chen, Kelvin Niu, George A. Komatsoulis, and Daoud Meerzaman. 2014. “OmicCircos: A Simple-to-Use R Package for the Circular Visualization of Multidimensional Omics Data.” Cancer Informatics 13: CIN–S13495.

Hubisz, Melissa J., Katherine S. Pollard, and Adam Siepel. 2011. “PHAST and RPHAST: Phylogenetic Analysis with Space/Time Models.” Briefings in Bioinformatics 12 (1): 41–51.

Jarvis, Erich D., Siavash Mirarab, Andre J. Aberer, Bo Li, Peter Houde, Cai Li, Simon YW Ho, Brant C. Faircloth, Benoit Nabholz, and Jason T. Howard. 2014. “Whole-Genome Analyses Resolve Early Branches in the Tree of Life of Modern Birds.” Science 346 (6215): 1320–1331.

Jurka, Jerzy, Vladimir V. Kapitonov, A. Pavlicek, P. Klonowski, O. Kohany, and J. Walichiewicz. 2005. “Repbase Update, a Database of Eukaryotic Repetitive Elements.” Cytogenetic and Genome Research 110 (1-4): 462–467.

Kapusta, Aurélie, and Alexander Suh. 2017. “Evolution of Bird Genomes—a Transposon’s-Eye View.” Ann NY Acad Sci 1389 (1): 164–185.

Katoh, Kazutaka, and Daron M. Standley. 2013. “MAFFT Multiple Sequence Alignment Software Version 7: Improvements in Performance and Usability.” Molecular Biology and Evolution 30 (4): 772–780.

Kaul, D., and H. A. Ansari. 1978. “Chromosome Studies in Three Species of Piciformes (Aves).” Genetica 48 (3): 193–96. https://doi.org/10.1007/BF00155569.

Koren, Sergey, Brian P. Walenz, Konstantin Berlin, Jason R. Miller, Nicholas H. Bergman, and Adam M. Phillippy. 2017. “Canu: Scalable and Accurate Long-Read Assembly via Adaptive k-Mer Weighting and Repeat Separation.” Genome Research 27 (5): 722–736.

Korf, Ian. 2004. “Gene Finding in Novel Genomes.” BMC Bioinformatics 5 (1): 59.

Korlach, Jonas, Gregory Gedman, Sarah B. Kingan, Chen-Shan Chin, Jason T. Howard, Jean-Nicolas Audet, Lindsey Cantin, and Erich D. Jarvis. 2017. “De Novo PacBio Long-Read and Phased Avian Genome Assemblies Correct and Add to Reference Genes Generated with Intermediate and Short Reads.” Gigascience 6 (10): gix085.

Kratochwil, Claudius F., and Axel Meyer. 2015. “Closing the Genotype–Phenotype Gap: Emerging Technologies for Evolutionary Genetics in Ecological Model Vertebrate Systems.” BioEssays 37 (2): 213–226.

Kurtz, Stefan, Adam Phillippy, Arthur L. Delcher, Michael Smoot, Martin Shumway, Corina Antonescu, and Steven L. Salzberg. 2004. “Versatile and Open Software for Comparing Large Genomes.” Genome Biology 5 (2): R12.

Li, Heng, and Richard Durbin. 2010. “Fast and Accurate Long-Read Alignment with Burrows– Wheeler Transform.” Bioinformatics 26 (5): 589–595.

Li, Heng, Bob Handsaker, Alec Wysoker, Tim Fennell, Jue Ruan, Nils Homer, Gabor Marth, Goncalo Abecasis, and Richard Durbin. 2009. “The Sequence Alignment/Map Format and SAMtools.” Bioinformatics 25 (16): 2078–2079.

Low, Wai Yee, Rick Tearle, Derek M. Bickhart, Benjamin D. Rosen, Sarah B. Kingan, Thomas Swale, Françoise Thibaud-Nissen, Terence D. Murphy, Rachel Young, and Lucas Lefevre. 2019. “Chromosome-Level Assembly of the Water Buffalo Genome Surpasses Human and Goat Genomes in Sequence Contiguity.” Nature Communications 10 (1): 1–11.

Manthey, Joseph D., Robert G. Moyle, and Stéphane Boissinot. 2018. “Multiple and Independent Phases of Transposable Element Amplification in the Genomes of Piciformes (Woodpeckers and Allies).” Genome Biology and Evolution 10 (6): 1445–1456.

Moore, William S., and Walter D. Koenig. 1986. “Comparative Reproductive Success of Yellow-Shafted, Red-Shafted, and Hybrid Flickers across a Hybrid Zone.” The Auk 103 (1): 42–51.

Nadachowska-Brzyska, Krystyna, Cai Li, Linnea Smeds, Guojie Zhang, and Hans Ellegren. 2015. “Temporal Dynamics of Avian Populations during Pleistocene Revealed by Whole-Genome Sequences.” Current Biology 25 (10): 1375–1380.

Notredame, Cédric, Desmond G. Higgins, and Jaap Heringa. 2000. “T-Coffee: A Novel Method for Fast and Accurate Multiple Sequence Alignment.” Journal of Molecular Biology 302 (1): 205–217.

Oliveira, Thays Duarte de, Rafael Kretschmer, Natasha Avila Bertocchi, Tiago Marafiga Degrandi, Edivaldo Herculano Corrêa de Oliveira, Marcelo de Bello Cioffi, Analía del Valle Garnero, and Ricardo José Gunski. 2017. “Genomic Organization of Repetitive DNA in Woodpeckers (Aves, Piciformes): Implications for Karyotype and ZW Sex Chromosome Differentiation.” Edited by Arthur J. Lustig. PLOS ONE 12 (1): e0169987. https://doi.org/10.1371/journal.pone.0169987.

Pagès, H., P. Aboyoun, R. Gentleman, and S. DebRoy. 2017. “Biostrings: Efficient Manipulation of Biological Strings.” R Package Version 2 (0).

Paradis, Emmanuel, Julien Claude, and Korbinian Strimmer. 2004. “APE: Analyses of Phylogenetics and Evolution in R Language.” Bioinformatics 20 (2): 289–290.

Platt, Roy N., Laura Blanco-Berdugo, and David A. Ray. 2016. “Accurate Transposable Element Annotation Is Vital When Analyzing New Genome Assemblies.” Genome Biology and Evolution 8 (2): 403–10. https://doi.org/10.1093/gbe/evw009.

Pollock, D. L., and N. S. Fechheimer. 1976. “The Chromosome Number of *Gallus Domesticus* 1.” British Poultry Science 17 (1): 39–42. https://doi.org/10.1080/00071667608416247.

Price, Alkes L., Neil C. Jones, and Pavel A. Pevzner. 2005. “De Novo Identification of Repeat Families in Large Genomes.” Bioinformatics 21 (suppl_1): i351–i358.

Quinlan, Aaron R., and Ira M. Hall. 2010. “BEDTools: A Flexible Suite of Utilities for Comparing Genomic Features.” Bioinformatics 26 (6): 841–842.

Seppey, Mathieu, Mosè Manni, and Evgeny M. Zdobnov. 2019. “BUSCO: Assessing Genome Assembly and Annotation Completeness.” In Gene Prediction, 227–245. Springer.

Short, L. L. 1982. “Woodpeckers of the World Greenville.” DE Delaware Museum of Natural History.

Simão, Felipe A., Robert M. Waterhouse, Panagiotis Ioannidis, Evgenia V. Kriventseva, and Evgeny M. Zdobnov. 2015. “BUSCO: Assessing Genome Assembly and Annotation Completeness with Single-Copy Orthologs.” Bioinformatics 31 (19): 3210–3212.

Smit, A., R. Hubley, and P. Green. 2015. “RepeatMasker Open-4.0. 2013-2015.” Institute for Sytems Biology. http://repeatmasker.org.

Smit, Arian FA, and Robert Hubley. 2008. “RepeatModeler Open-1.0.” Available Fom Http://Www.Repeatmasker.Org.

Stanke, Mario, Mark Diekhans, Robert Baertsch, and David Haussler. 2008. “Using Native and Syntenically Mapped CDNA Alignments to Improve de Novo Gene Finding.” Bioinformatics 24 (5): 637–644.

Team, R. Core. 2018. “R: A Language and Environment for Statistical Computing.[Google Scholar].”

Toews, David P.L., Scott A. Taylor, Rachel Vallender, Alan Brelsford, Bronwyn G. Butcher, Philipp W. Messer, and Irby J. Lovette. 2016. “Plumage Genes and Little Else Distinguish the Genomes of Hybridizing Warblers.” Current Biology 26 (17): 2313–18. https://doi.org/10.1016/j.cub.2016.06.034.

Walker, Bruce J., Thomas Abeel, Terrance Shea, Margaret Priest, Amr Abouelliel, Sharadha Sakthikumar, Christina A. Cuomo, Qiandong Zeng, Jennifer Wortman, and Sarah K. Young. 2014. “Pilon: An Integrated Tool for Comprehensive Microbial Variant Detection and Genome Assembly Improvement.” PloS One 9 (11): e112963.

Warren, René L., Chen Yang, Benjamin P. Vandervalk, Bahar Behsaz, Albert Lagman, Steven JM Jones, and Inanç Birol. 2015. “LINKS: Scalable, Alignment-Free Scaffolding of Draft Genomes with Long Reads.” GigaScience 4 (1): s13742–015.

Waterhouse, Robert M., Mathieu Seppey, Felipe A. Simão, Mosè Manni, Panagiotis Ioannidis, Guennadi Klioutchnikov, Evgenia V. Kriventseva, and Evgeny M. Zdobnov. 2018. “BUSCO Applications from Quality Assessments to Gene Prediction and Phylogenomics.” Molecular Biology and Evolution 35 (3): 543–548.

Wiebe, K. L. 2020. “Northern Flicker (Colaptes Auratus), Version 1.0. In Birds of the World(P.G. Rodewald, Editor).” 2020. https://doi.org/10.2173/bow.norfli.01.

Wiebe, Karen L. 2000. “Assortative Mating by Color in a Population of Hybrid Northern Flickers.” The Auk 117 (2): 525–529.

Wiley, Graham, and Matthew J. Miller. 2020. “A Highly Contiguous Genome for the Golden-Fronted Woodpecker (Melanerpes Aurifrons) via Hybrid Oxford Nanopore and Short Read Assembly.” G3: Genes, Genomes, Genetics 10 (6): 1829–1836.

Wright, Natalie A., T. Ryan Gregory, and Christopher C. Witt. 2014. “Metabolic ‘Engines’ of Flight Drive Genome Size Reduction in Birds.” Proceedings of the Royal Society B: Biological Sciences 281 (1779): 20132780. https://doi.org/10.1098/rspb.2013.2780.

Yang, Ziheng. 1997. “PAML: A Program Package for Phylogenetic Analysis by Maximum Likelihood.” Computer Applications in the Biosciences 13 (5): 555–556.

Yeo, Sarah, Lauren Coombe, René L. Warren, Justin Chu, and Inanç Birol. 2018. “ARCS: Scaffolding Genome Drafts with Linked Reads.” Bioinformatics 34 (5): 725–731.

Zhang, Guojie, Cai Li, Qiye Li, Bo Li, Denis M. Larkin, Chul Lee, Jay F. Storz, Agostinho Antunes, Matthew J. Greenwold, and Robert W. Meredith. 2014. “Comparative Genomics Reveals Insights into Avian Genome Evolution and Adaptation.” Science 346 (6215): 1311–1320.

